# Object manifold geometry across the mouse cortical visual hierarchy

**DOI:** 10.1101/2020.08.20.258798

**Authors:** Emmanouil Froudarakis, Uri Cohen, Maria Diamantaki, Saumil Patel, Zheng Tan, Taliah Muhammad, Edgar Y. Walker, Jacob Reimer, Philipp Berens, Haim Sompolinsky, Andreas S. Tolias

## Abstract

Despite variations in appearance, we recognize objects robustly. Neuronal populations responding to objects presented under varying transformations form object manifolds, and hierarchically organized visual areas are hypothesized to untangle pixel intensities into linearly decodable object representations. However, the associated changes in the geometry of the object manifolds along the cortex remain unknown. Using home cage training, we showed that mice are capable of invariant object recognition. We recorded the responses of thousands of neurons to measure the information about object identity across the visual cortex and found that the lateral areas LM, LI, and AL carry more linearly decodable object information compared to other visual areas. We applied the theory of linear separability of manifolds and found that the increase in classification capacity is associated with a decrease in the dimension and radius of the object manifold, identifying the key features in the geometry of the population neural code that are associated with invariant object coding.

## Introduction

Object recognition is an ethologically-relevant task for many animals. This is a challenging problem because an individual object can elicit myriads of images on the retina due to so-called nuisance transformations such as changes in viewing distance, projection, occlusion, and illumination. The collection of neural population responses associated with a single object is known as the object manifold. A prevailing hypothesis is that along the visual hierarchy, object manifolds are gradually untangled to produce increasingly invariant object representations, which are linearly decodable (Dicarlo and Cox, 2007). This hypothesis is primarily based on work in non-human primates, which is a powerful model to study object recognition, especially given the similarities in visual perception among primates. These studies have revealed that the selectivity for object identity increases as visual signals are conveyed from primary visual cortex (V1) to inferotemporal cortex (Hung et al., 2005; Rust and Dicarlo, 2010). Despite this significant progress, the underlying changes in the geometry of the object manifolds along the visual cortical hierarchy that lead to object recognition remain largely unknown. The mouse animal model is ideally suited to dissect circuit mechanisms due to its genetic tractability and the numerous methods available to perform large-scale recordings, manipulations, and anatomical tracing with cell-type precision (Fenno et al., 2015; Sofroniew et al., 2016). Therefore developing visually guided behaviors in rodents is important (Zoccolan et al., 2009) and identifying the relevant network of visual areas involved in object recognition analogous to the ventral stream of primates is critical towards an eventually circuit level mechanistic dissection of the algorithms of object recognition. In this direction, we developed a novel automatic high-throughput training paradigm and demonstrated that mice can be trained to perform a two-alternative forced choice (2AFC) object classification task, which is typically used in primates to test object identification. While visually-guided operant behavioral tasks have been used previously in mice (Han et al., 2022; Procacci et al., 2020; Leger et al., 2013), here we show that mice can also learn to correctly discriminate objects under a 2AFC paradigm. Critically, this capability persisted even when they were presented with previously unseen transformation of objects demonstrating that mice are capable of invariant object recognition.

To systematically study how objects are encoded in the mouse visual system, we recorded the activity of thousands of neurons across all cortical visual areas of the mouse: primary (V1), anterolateral (AL), rostrolateral (RL), lateromedial (LM), lateral intermediate (LI), posteromedial (PM), anteromedial (AM), posterior (P), postrhinal (POR) and laterolateral anterior (LLA) visual areas, while presenting images of moving objects undergoing numerous identity-preserving transformations such as rotation, scale and translation across different illumination conditions. By decoding the identity of the objects from the recorded neural activity using a linear classifier, we found that the lateral extrastriate visual areas (LM, AL, LI) carried more linearly decodable information about object identity compared to V1 and all the other higher-order areas we studied. The large scale recordings provide the opportunity to study the changes in the geometry of object manifolds along the cortex associated with invariant object coding. We applied the recently developed theory of linear separability of manifolds (Cohen et al., 2019; Chung et al., 2018) to our neural recordings and found that in areas LM and AL the increase in classification capacity is associated with distinct optimizations of the manifold geometry, where both the manifold radii and dimensions are reduced compared to other visual areas. Additionally, by recording simultaneously from many visual areas, we found that the population dynamics differed across the visual hierarchy, with information about object identity accumulating faster in areas that were more object selective compared to V1.

## Results

### Mice are capable of invariant object recognition

We generated movies of 3D objects by varying their location, scale, 3D pose and illumination in a continuous manner across time (**Fig. 1a, Supp. Movie 1**). We developed a 2AFC automatic home cage training system in which water restricted mice had to lick a left or a right port depending on the object that was shown on a small monitor in front of their cage (**Fig. 1b, Materials and Methods**). Upon a correct choice, animals immediately received a small amount of water reward. Naive animals initially licked the left and right probes at random, but within two weeks of training, animals learned to preferentially lick the correct port matched to object identity (**Fig. 1c**); trained animals maintained consistent performance on the task across days (**Fig. 1d**). An important property of object recognition is the ability to generalize across views of objects that have never been seen before. After the animals learned to discriminate objects from the movie clips - which contained a specific set of object transformations, new movie clips with unique parameters across translation, scale, pose and illumination were presented to the animals. We did not detect any differences in performance between the previously seen object transformations (**Fig. 1e**, familiar transformations) and novel object transformations (**Fig. 1e**, novel transformations). The ability to generalize across identity-preserving transformations indicated that mice learned an internal object-based model and did not rely simply on low-level features of the rendered movies they observed during training. Importantly, a linear classifier trained on the pixel intensities of the rendered movies performed at chance level (**Fig. 1e**, pixel model), indicating that the discrimination of these objects cannot be simply solved using low-level strategies based on pixel intensity differences between the images. Moreover, a model of the center-surround receptive fields (Difference of Guassians, DoG) was also lower than the behavioral performance, indicating the necessity of cortical computations (**Fig. 1e**, DoG model). To identify how different objects affect the behavioral performance, we first compared the performance across all four objects that were shown to the animals during the task (**Fig. 1f**). Behavioral discriminability was significantly better for Object B compared to objects A and D (Kruskal-Wallis, p<0.05). We then used a linear regression model to identify the latent parameters of the objects that contribute most to the behavioral variability. Object size had a significant positive effect on behavioral performance (i.e. larger objects were easier to discriminate) while position on the vertical axis (Y translation) had a small but significant negative effect (i.e. larger vertical offsets from the center were more difficult to discriminate, **Fig. 1g, Supp. Fig. 1**). Our behavioral results show that mice can discriminate between objects and are sensitive to specific visual features.

**Fig. 1.**
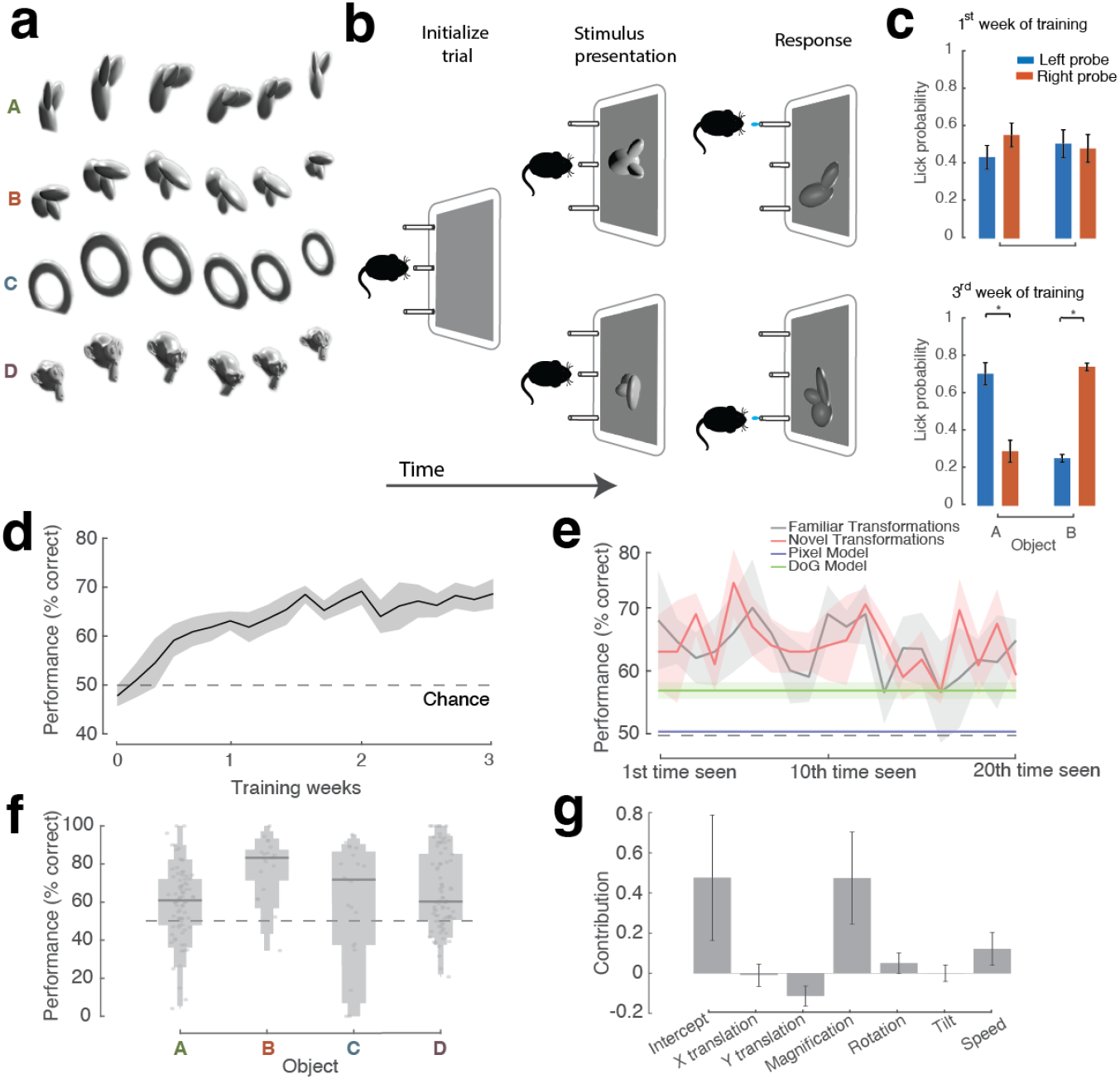
Experimental procedure for behavioral training. **a**. Single frames from movies with the objects that were presented to the animals. **b**. Behavioral training sequence. **c**. Probability of licking either probe during the early training period (upper bar plot) and later training period (lower bar plot) for 1 animal. Error bars represent S.E.M. Student t-test * p < 0.05 **d**. Performance as a function of training time, N = 7 animals. **e**. Performance across repetitions of familiar (gray) and novel (red) object transformations during one session. N = 5 animals. Gray dashed line represents chance level. Blue line represents performance of a pixel based linear classifier. G reen l ine represents performance of a DoG filter based linear classifier. F or both (d) and (e) shaded areas represent S.E.M. **f**. Performance for different objects. Lines indicate the median of the distribution, and the variable bars indicate the 75, 90, 95, and 100 percentiles. Dots represent the raw values. N = 5 animals. N= 80, 20, 20, 80 conditions for objects A, B, C and D respectively **g**. Contributions of the object latent parameters toward predicting the average performance of the animals. N = 5 animals. Error bars show s.e. of the regression coefficients scaled by the mean difference.

### Lateral visual areas carry more linearly decodable object identity information

If mice are capable of discriminating between objects, there should exist a set of areas along their visual processing hierarchy that can extract this information. It has been suggested that one way of extracting the object information irrespective of its transformations is to have neural representations for each object that are untangled, i.e. can be read-out using a linear decoder (Dicarlo and Cox, 2007). To test this idea, we used transgenic mice expressing GCaMP6s in pyramidal neurons and recorded the activity of hundreds of neurons in each visual area separately or of thousands of neurons across the whole visual cortical hierarchy of the mouse using a large field of view microscope ((Sofroniew et al., 2016), **Fig. 2a,b**), while the animals passively viewed the moving objects (**Fig. 1a**). We identified the borders between visual areas using wide-field retinotopic mapping as previously described (Fahey et al., 2019; Garrett et al., 2014; Wang and Burkhalter, 2007) (**Fig. 2a, Materials and Methods**). In total we recorded 489182 neurons: 222239 in V1, 58159 in RL, 48769 in LM, 24662 in AL, 16681 in PM, 15364 in AM, 11617 in LI, 6845 in LLA, 6231 in P, 6029 in POR, and 72586 neurons not assigned to any visual area. Neurons in all identified visual areas showed significantly more reliable responses compared to neurons that were not assigned to any visual area (**Supp. Fig. 2a**).

**Fig. 2.**
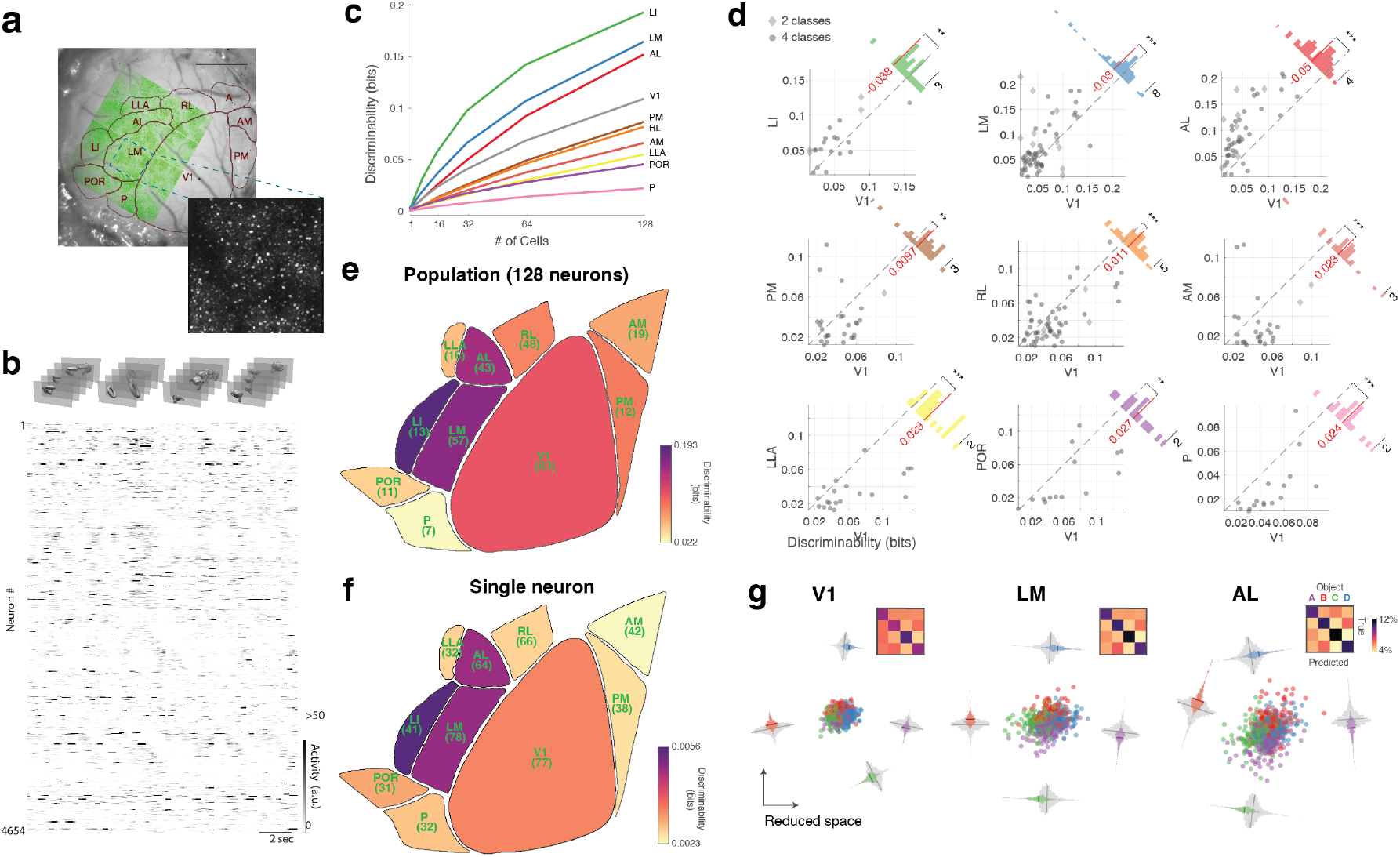
Object identity decoding across the visual hierarchy. **a**. Example large field of view recording (green) with area boundaries overlaid. Scale bar represents 1mm. A small inset depicts the two-photon average image for a small segment of the large field of view captured with the mesoscope. **b**. Example responses of all neurons to moving objects (shown on top) from the recording shown in (a). Each clip is presented for 3-5 seconds before a short pause switches to a new clip that might be the same or a different object identity. **c**. Discriminability of object identity as a function of the number of neurons sampled. Each line represents the average across all recorded sites. **d**. Scatter plot of the discriminability of different areas with a population of 64 neurons compared to V1 for all the recording sites. Insert histogram represents the difference between the discriminability of each area and V1. Red line and number indicate the mean difference. Diamonds represent the results with 2 objects whereas circles represent the results with 4 objects. Outliers have been omitted for better visualization. Wilcoxon signed rank test *** p < 0.001, ** p <0.01, * p < 0.05. **e**. Average discriminability of all visual areas with a population of 128 neurons. The number below each area represents the recording sites sampled. **f**. Same as in (e) but when using a single neuron at a time to decode the object identity. The number below each area represents the cells sampled. **g**. Low-dimensional representation of the 128-dimensional neural activity space, illustrating the separation of the responses to four different objects for three example areas. Each dot represents the average of the activity in one 500msec bin. The side histograms represent the distances of the data projected onto each of the four object category axes for the same-class (colored) and different-class (gray). Each insert represents the confusion matrix after decoding.

To measure how linearly discriminable the responses to the different objects were, we used cross-validated logistic regression to classify the object identity from the responses of neurons in each visual area. As expected, discriminability increased as a function of the number of neurons sampled (**Fig. 1c, Supp. Fig. 3c**), but only the higher visual areas, LM, LI, and AL, showed consistently higher discriminability levels compared to V1 responses (**Fig. 2c,e**) for a given number of neurons. In contrast, areas RL, PM, AM, P, POR, and LLA had significantly lower discriminability levels compared to V1 and this effect was independent of the number of objects (**Fig. 2d**). The differences in decoding across areas persisted at the single neuron level as well (**Fig. 2f, Supp. Fig. 3a**).

**Fig. 3.**
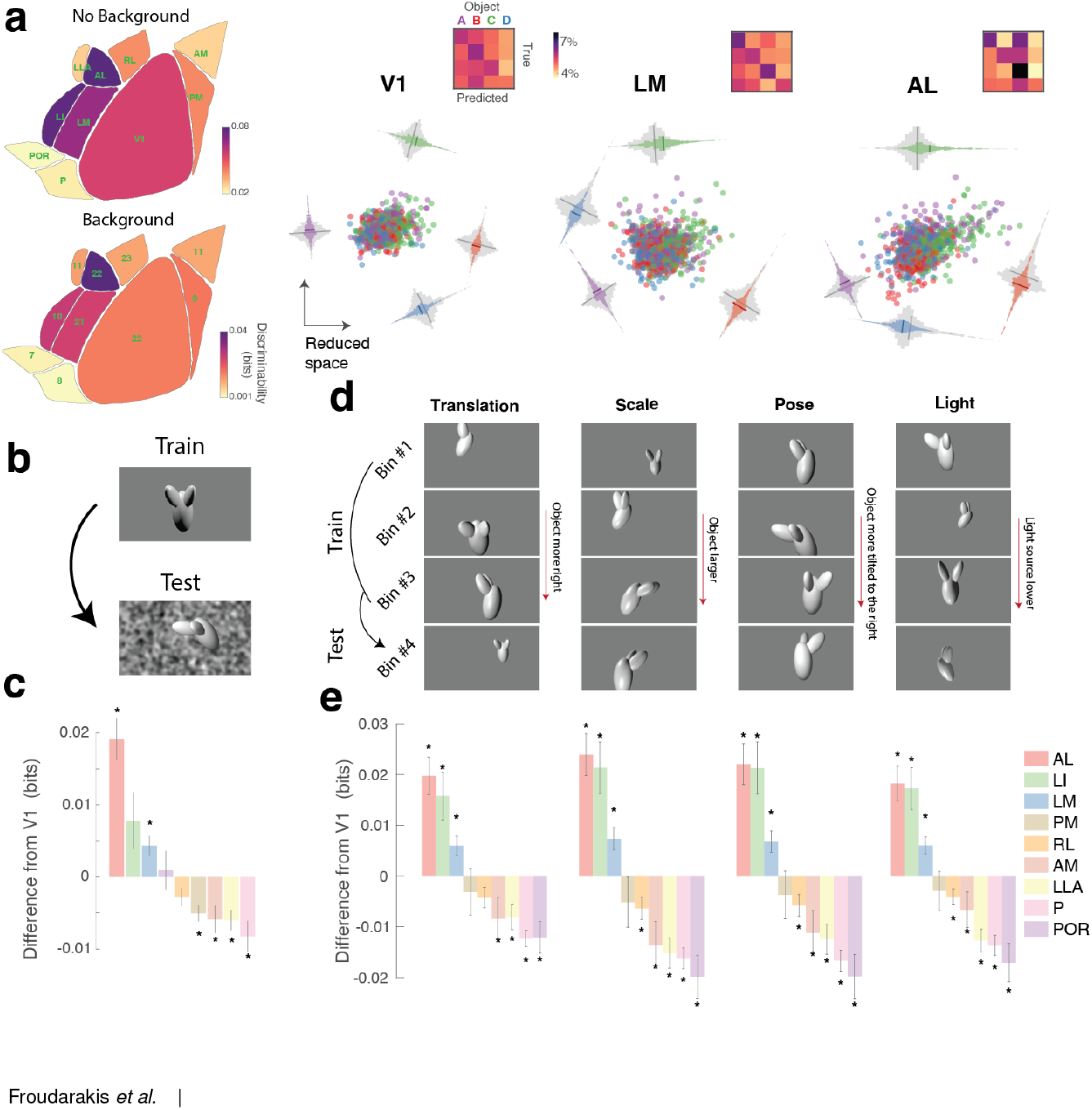
Generalization performance across background noise and identity-preserving transformations. **a**. Left Top: Average performance of the decoder for objects without background. Left Bottom: Average performance of the decoder for objects with background. Right:Low-dimensional representation of the responses to the object w/ background are shown on the right similar to Fig. 2e. Each insert represents the confusion matrix after decoding. **b**. Generalization test across background noise. The decoder was trained on the responses to objects without background and tested on the responses to objects that contained background noise. **c**. Bar plot indicating the difference in performance when testing the ability of the decoder to generalize across background noise, compared to the performance of the decoder in V1. * indicate p < 0.05 Kruskal-Wallis with multiple comparisons test. **d**. Example parameter space of the four nuisance classes: Translation (x/y), Scale, Pose (tilt/rotation) and Light (four light sources). The example images for the 4 bins are depicting objects that are located more to the right (Translation), objects that are larger (Scale), objects that are rotated more to the right (Pose) and a light source that is located lower (Light). The decoder was tested on a parameter space of each of the four nuisance variables that had not been part of the training set. **e**. Bar plot indicating the difference of the performance when testing on untrained parameter space, compared to the performance of V1. * indicate p < 0.05 Kruskal-Wallis with multiple comparisons test.

We performed several control analyses: As previously reported (Garrett et al., 2014), the coverage of the visual field is different between visual areas. To control for our results being affected by differences in the retinotopic coverage across areas, in some experiments we mapped the receptive fields (RF) of all recorded neurons using a dot stimulus (see **Materials and Methods**) and repeated our decoding analysis using only neurons from each area with RF centers within the same ~20° area of visual space. Areas LM, LI, and AL still showed significantly higher discriminability (**Supp. Fig. 3b**). When we estimated the decoding performance for pairwise object discrimination using the total number of neurons sampled from each area, we found that several areas (V1, LM, RL and AL) showed decoding performance that equaled or surpassed the behavioral performance, indicating that these areas contain sufficient information to explain the observed behavior (**Supp. Fig. 3c,d**).

Another potential confound might be differences in RF sizes across areas (Wang and Burkhalter, 2007; Murgas et al., 2020). To investigate the influence of RF size on discriminability, we modeled the effect of changing RF size in a simulated population of neurons using the output of Gabor-like filters learned by a sparse coding model of natural images using independent component analysis (ICA) (van Hateren and van der Schaaf, 1998). ICA filters (**Supp. Fig. 4a**) resulted in a much higher performance compared to a simple pixel-based model (**Supp. Fig. 4b**), but still almost an order of magnitude less than the neural data (Compare **Supp. Fig. 4b** Random to **Fig. 2c**). Increasing the size of the RF by either scaling or pairwise linearly combining them (**Supp. Fig. 4a**, ICA 150% and ICA multi respectively, see **Materials and Methods**) led to either a decrease in discriminability or had no significant effect, respectively (**Supp. Fig. 4b**). These results argue that our *in vivo* results cannot be explained by differences in the RF sizes across visual areas.

**Fig. 4.**
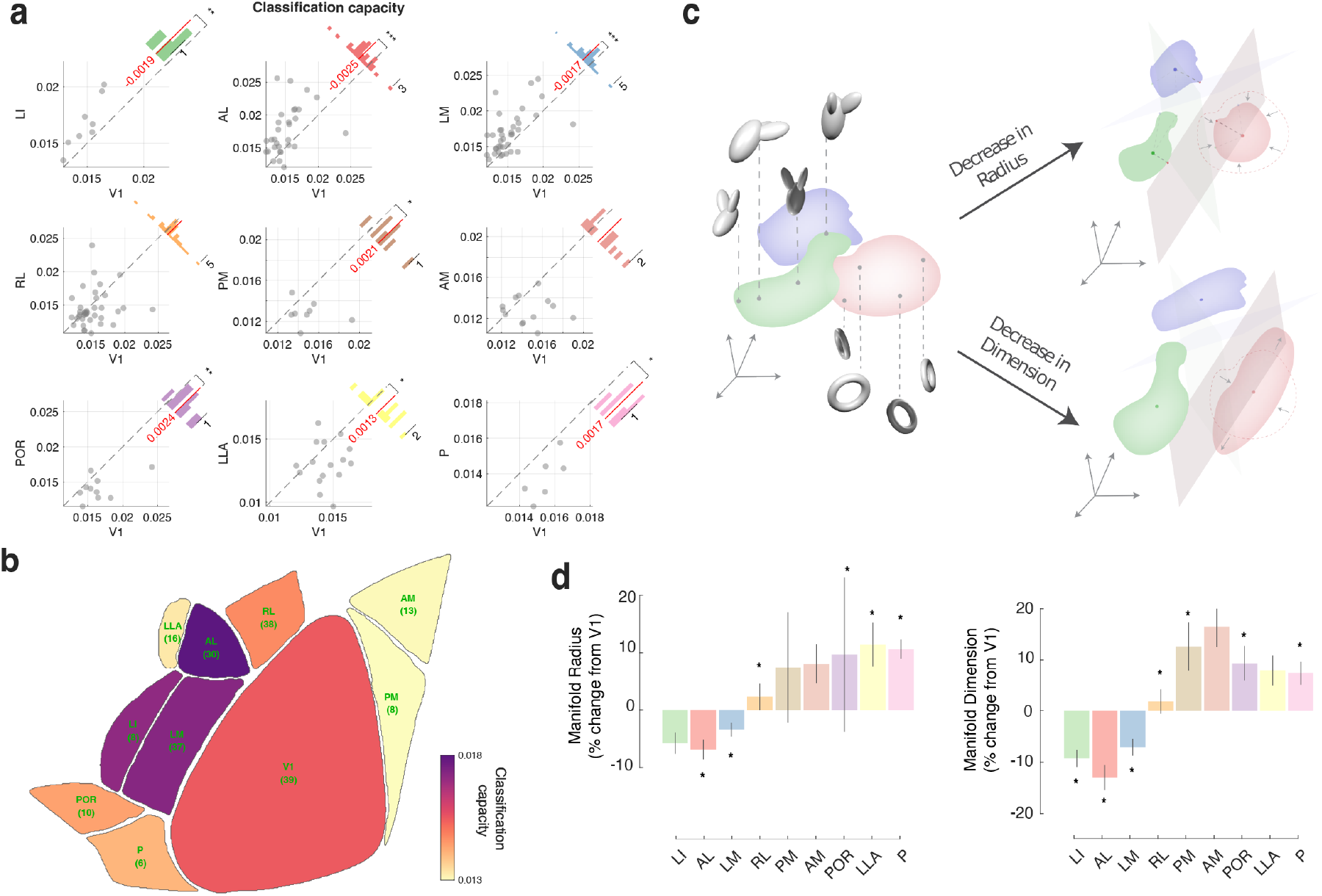
Classification capacity and geometry of manifolds across the visual hierarchy. **a**. Scatter plot of the classification capacity of different areas compared to V1 for 4 objects. Insert histogram represents the difference between the classification capacity of each area and V1. Red line and number indicate the mean difference. Wilcoxon signed rank test *** p < 0.001, ** p < 0.01, * p < 0.05. **b**. Average classification capacity of all visual areas with a population of 128 neurons. The number below each area represents the recording sites sampled. **c**. Illustration of low dimensional representations of object manifolds for two visual areas. Left: each point in an object manifold corresponds to neural responses to an object under certain identity-preserving transformations. Right: demonstration of two possible changes in the manifold geometry in a higher order area, reduction of the radius of one manifold through reduction of its extent in all directions (top) and reduction of the dimension of one manifold by concentrating variability at certain elongated axis, reducing the spread along other axes. Such changes have predictable effects on the ability to perform linear classification of those objects. **d**. Bar plots of the manifold radius differences to V1 (left), and manifold dimension differences to V1 (right) of all areas. Wilcoxon signed rank test, * p < 0.05.

An area with larger RF might be better at representing objects simply because more neurons respond to the object at any moment. Indeed, when we examined decoding performance conditioned on the object size, we observed an increase in discriminability for all visual areas as a function of object size (**Supp. Fig. 5a**), in agreement with the increased performance we found when sampling from more neurons (**Fig. 2c**) and the effect on behavioral performance (**Fig. 1g**). However, if changes in RF size alone are responsible for increased object discriminability, we would expect that area PM, which has very large RF (Murgas et al., 2020), would also have high object discriminability. This was not the case in our data (**Fig. 2e,d**). Another object parameter that we found affects behavioral performance was speed (**Fig. 1g**) and higher visual areas have been reported to have different temporal frequency selectivities (Han et al., 2022; Andermann et al., 2011; Marshel et al., 2011). To determine whether differences in speed selectivity could explain our results, we analyzed decoding performance as a function of object speed. Unlike the strong effect we found for object size (**Supp. Fig. 5a**), speed had minimal impact on decoding performance (**Supp. Fig. 5b**), with only minor differences between areas LI, PM and AM at very low and very high speeds (**Supp. Fig. 5c**).

**Fig. 5.**
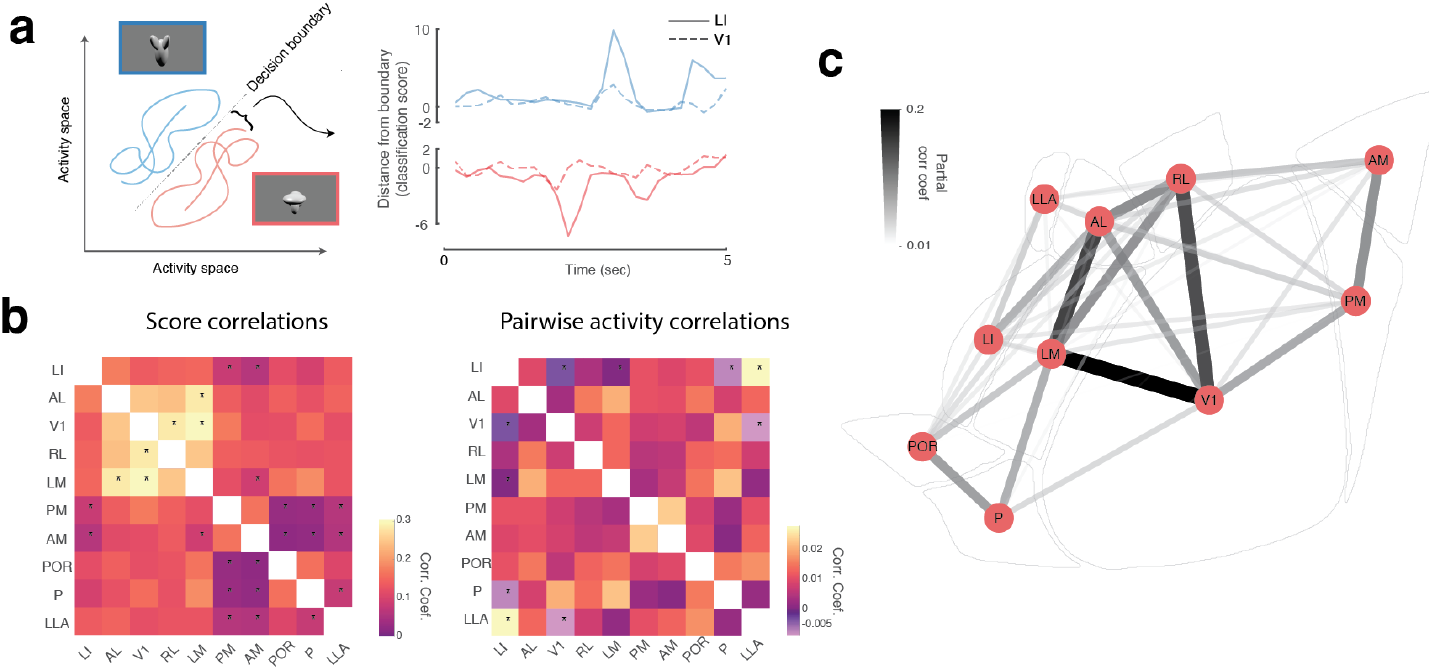
Temporal dynamics and cross-area dependencies. **a**. Schematic representation of the classification scores as the distances of the response trajectories to the decision boundary (left) and their resulting temporal dependencies across different areas (right). **b**. Score correlations across all recorded areas (left) and raw pairwise correlations of the single neuron activity between areas (right). Significance was estimated by bootstrapping across all correlations, * p < 0.025/45. **c**. Schematic representation of the score partial correlation coefficients between areas.

Finally, we have previously shown that the animal’s behavioral state can affect neural responses to visual stimuli (Froudarakis et al., 2014; Reimer et al., 2014; Froudarakis et al., 2019; McGinley et al., 2015). To determine whether our decoding results are affected by the animal’s behavior, we split the data into periods where the animals were running and periods that the animals were stationary. Although there was an increase in decoding performance when the animals were running, the effect was apparent for all visual cortex and areas LI, LM and AL had still significantly higher performance than the rest (**Supp. Fig. 6**, Running vs Stationary).

Taking into account the results from **Fig. 2** and the afore-mentioned controls, the increase in discriminability in AL, LI, and LM cannot be explained by low level RF, object or behavioral state properties, instead these data indicate that these visual areas are particularly involved in the processing of visual object information and thus their neural representations of objects are easier to separate (**Fig. 2g**).

### Lateral visual areas show responses that are more invariant to nuisance transformations

An important property of visual areas that extract information about object identity is generalization to out-of-distribution data, such as adding background noise to the stimuli. To assess the effect of background noise, we first trained a logistic regression decoder on responses to objects with only a gray background as previously used. We then evaluated the performance of the decoder on the responses to movies in which we embedded the objects on top of background noise (**Fig. 3b, Supp. Movie 2**). While the discriminability decreased for all visual areas when compared to noise-free stimuli (**Fig. 3a**), area AL maintained significantly higher discriminability compared to V1 (**Fig. 3a,c**), indicating that in addition to being highly invariant to changes in the appearance of the object, the object representation in this area is also more robust than in V1 and other visual areas to background noise. The drop in discriminability cannot be explained by a drop in reliability, as adding noise did not affect the reliability of the neural responses (**Supp. Fig. 2b**).

An additional test of invariant object recognition is the ability of the neural representation to generalize across new image transformations — not used during training — that preserve object identity such as changes in position, scale, pose and illumination (Dicarlo and Cox, 2007; Rust and Dicarlo, 2010; Henaff et al., 2019; Tafazoli et al., 2017). Specifically an object decoder built on a subset of the nuisance parameter space, i.e. a limited range of translations, sizes, and rotations, should generalize across new nuisance parameters. To test this we split the data into four non-overlapping bins for each of the nine continuously-varying parameters that defined the object stimulus (for example for the size object parameter into very small, small, medium, and large objects; **Fig. 3d**), while the remaining parameters were randomly sampled. For each parameter, we then used data from three of the bins to train the decoder and tested the prediction performance on the held-out data bin (4-fold cross validation). Comparing the out-of-distribution test set performance of each area to the performance of V1 allowed us to characterize the invariance of the representation in each area in relation to V1 (**Fig. 3e**). Areas AL, LM, and LI consistently showed the best generalization performance, when changing translation, scale, pose or light (**Fig. 3e**). The larger RF sizes of areas PM and AM (Wang and Burkhalter, 2007; Murgas et al., 2020) do not seem to contribute to an improved translation invariance.

### Changes in the geometry of object manifolds along the cortical hierarchy

Chung and colleagues (Chung et al., 2018) recently developed the theory of linear separability of manifolds and defined a measure called the classification capacity which quantifies how well a neural population supports object classification. The classification capacity measures the ratio between the number of objects and the size of the neuronal population that is required for reliable binary classification of the objects, and is tightly related to the geometry of a neuronal population responding to an object presented under varying nuisance transformations with respect to the identity of the object (object manifold). In deep neural networks trained on object classification tasks, it has been shown that the classification capacity improves along the network’s processing stages (Cohen et al., 2019). Our data, consisting of responses of large neuronal populations in different visual areas to objects under various transformations, are well suited for applying this method to characterize the object manifolds in different visual areas. We used the neuronal responses of 128 simultaneously recorded neurons from each visual area to four objects under the identity-preserving transformations introduced earlier (object position, scale, pose and illumination conditions, with and without background noise). In agreement with our decoding results, we found that the classification capacity increased in higher visual areas LI, AL and LM compared to V1, but decreased in the rest of the areas (**Fig. 4a,b**). The estimated capacity was significantly higher compared to the control case, were the object identity was shuffled across trials (**Supp. Fig. 7**). The theory of linear separability of manifolds (Chung et al., 2018) also enabled us to characterize the associated changes in the geometry of the object manifolds to understand how object invariant representations arise along the processing hierarchy (Cohen et al., 2019) (i.e. relate the manifolds’ classification ability to the geometry of object manifolds). In particular, classification capacity depends on the overall extent of variability across the encoding dimensions, the radius of the manifold, but also the number of directions in which this variability is spread, the dimension of the manifold. These geometric measures influence the ability to linearly separate the manifolds (**Fig. 4c**). In our results, we find that the increase in classification capacity can be traced to changes in the manifolds’ geometry, both as a decrease of the dimension and radius of object manifolds (**Fig. 4d**).

### Temporal dynamics and cross-area dependencies

One question that arises is how these visual areas are able to form invariant representations that can generalize across background noise or nuisance parameters. One way for these areas to optimize the representations is by taking advantage of the temporal continuity that exists for natural objects by integrating information over time (Dicarlo et al., 2012; Orlov and Zohary, 2017). We analyzed the temporal dynamics of the decoding performance of random samples of 50 simultaneously recorded neurons for objects overlaid on background noise. From one trial to the next the nuisance parameters varied continuously but the object identity was preserved (cis trials) or switched (trans trials) (**Fig. 2b**). When we compared the discriminability as a function of time for cis/trans trials, we found that indeed in the trials in which the identity of the object was switched (trans trials), discriminability was overall lower across all visual areas in the early phase of the trials compared to the late phase of the trials, providing evidence for temporal integration during a trial (**Supp. Fig. 8a,b**). During the 0.5-1.5 seconds of the trans trials discriminability in several areas was significantly higher than V1, suggesting that object identity is represented in these higher visual areas earlier than in V1 (**Supp. Fig. 8c,d**).

We also studied the correlations between the representations of objects across multiple visual areas. If information about object identity propagates across areas, then we expect to find significant temporal correlations in the evolution of object discriminability across these areas. We estimated each area’s confidence about the identity of the object at each time point within the trial for the class that was presented, as the distance of the population activity from the decision boundary (**Fig. 5a**), and we examined the co-fluctuation of this metric across time across the visual areas by computing the correlation between these vectors (Score Correlation).The highest correlations in this moment-to-moment discriminability score were between AL, LM, RL and V1 (**Fig. 5b (left), c**).

Given that activities of neurons across areas can co-fluctuate because of global brain states, these score correlations could just be the result of raw activity correlations across areas. To test this we computed the activity correlations between the responses of pairs of neurons across visual areas(**Fig. 5b (right)**). We observed a different correlation pattern that was distinct from the structure of the score correlation (**Fig. 5b**). Moreover, we measured the strength of the linear relationship between each pair of areas after adjusting for relationships with the rest of the areas. To this end, we computed the partial score correlations. The correlation pattern remained largely unchanged with strong dependencies between V1-LM, V1-RL and LM-AL (**Fig. 5c**) suggesting that these areas work together as a network of areas specialized for object recognition. Interestingly, we did not find a strong relationship between V1-AL (**Fig. 5c**). In order to evaluate how these cross-area dependencies are influenced by different object latent parameters we built a decoder that can separate the latent parameter as well as the object identity (see **Materials and Methods**). Adding the background noise had a significant increase of correlations across the majority of the areas, and a similar effect was observed but to a lesser extent with the translation and scale latent parameters (**Supp. Fig. 9a,b**). Interestingly, area POR showed increased score correlations with almost all the other areas when also decoding object translation (**Supp. Fig. 9b Purple**) which has been shown to receive input from superior colliculus (Beltramo and Scanziani, 2019).

## Discussion

The ability to recognize, discriminate, and track objects across time is a key adaptive trait that is fundamental to identifying food items or conspecifics (Jones and Ratterman, 2009). The ability to recognize objects has been observed not only in higher mammals such as humans and monkeys, but also rodents, birds, fish and insects (Zoccolan et al., 2009; Bevins and Besheer, 2006; Blaser and Heyser, 2015; Newport et al., 2018; Soto and Wasserman, 2014; Werner et al., 2016). While the implementation of how object information is extracted from the visual scene may vary across species, the computational problem remains the same: construct an invariant representation of objects under a wide range of identity-preserving transformations.

In this work, we showed that mice can be trained to recognize unfamiliar objects in a 2AFC paradigm (**Fig. 1**). While there is plenty of evidence that mice can detect novel objects (Leger et al., 2013), and that mice rely on their vision to hunt crickets (Hoy et al., 2016), until our study there was no direct evidence that mice are capable of invariant object recognition. Similar tasks have been developed for rats (Zoccolan et al., 2009), but mice have not been reported to perform such a task. That might be related to the fact that even though mice and rats can achieve similar performance levels, mice are slower to train (Jaramillo and Zador, 2014). Our unique training approach involves minimal interactions with the animals since the training system is part of their housing. Within a few weeks animals learn to discriminate objects and can show generalization across unseen objects poses and background noise establishing that mice are capable of invariant object recognition (**Fig. 1d**).

To identify how animals are able to extract object identity, we analyzed the activity of thousands of neurons of all known visual cortical areas of the mouse. We found that the decoding performance varied across the visual hierarchy where a set of lateral visual areas carried more linearly decodable information about the object identity. Importantly, these areas retained the information about object identity even in difficult visual conditions such as background noise and across new identity-preserving transformations despite that the linear classifiers were not trained under those conditions. Our results agree with the hypothesis that object representations become untangled and more linearly separable as information progresses through the visual hierarchy (Dicarlo et al., 2012; Lindsey and Issa, 2024). This process might be beneficial, as a simple read-out mechanism can be employed to drive behavior. It is important to note that a biologically plausible readout mechanism could involve only from a small set of projection neurons in order to extract object identity. We found that information carried by single neurons also increased progressively across the hierarchy of V1-LM-LI in lateral visual cortex in agreement with electrophysiology studies in the rat (Tafazoli et al., 2017) (**Supp. Fig. 3a**). Importantly, our richly varied object stimuli cannot be easily discriminated by a simple pixel or DoG model (**Fig. 1b, Supp. Fig. 5b,c**) and the ability of mice to separate such objects likely depends on more complex computations in cortical circuits. Recent electrophysiological work in macaques confirms that classification capacity and object identity factorization improve from V1 through V4 to IT cortex, supporting a shared representational strategy across species (Cadena et al., 2024). Therefore, analogous to primates, hierarchically organized visual areas in mice untangle pixel intensities into more linearly decodable object representations.

However, the associated changes in the geometry of the object manifolds along the visual cortex remained unknown. To this end, we characterized how the geometry of the object manifolds changed across the visual hierarchy, using the newly developed theory of linear separability of manifolds (Cohen et al., 2019; Chung et al., 2018). We found that three lateral visual areas LI, LM and AL showed increased classification capacity with object manifolds becoming smaller and having lower dimensionality (**Fig. 4**). While the classification capacity and radius of object manifolds has not been previously quantified along the visual processing hierarchy, our results on the dimensionality of the neural population agree with previous work. Different methods have been used to quantify the dimensionality of the population responses which also showed that it decreases along the visual hierarchy of monkeys (Brincat et al., 2018; Lehky et al., 2014). However, critically the theory of the linear separability of manifolds differs from these previous methods as it quantifies the geometrical properties of the object response manifolds which contribute to the ability to perform linear decoding. This enabled us to determine that the dimension of the object manifold decreases from primary visual cortex to higher visual areas in a way which allows for linear decoding of objects using smaller number of neurons.

The higher visual areas of the mouse (Wang and Burkhalter, 2007; Glickfeld and Olsen, 2017), have distinct spatiotemporal selectivities (Andermann et al., 2011; Marshel et al., 2011) and project to different targets (Wang et al., 2012). Based on these differences in their selectivities, projection and chemoarchitectonic patterns, efforts have been made to separate areas into ventral and dorsal pathways analogous to those described in primates (Wang et al., 2012; Smith et al., 2017; Wang et al., 2011; Wang and Burkhalter, 2013). Specifically, areas such as LM, LI, P and POR areas are hypothesized to comprise the ventral stream whereas areas AL, RL, AM and PM comprise the dorsal stream. In rats, lateral visual areas LM, LI and LL have been shown to carry progressively more information about objects (Tafazoli et al., 2017; Vermaercke et al., 2015, 2014). However, the areas of the mouse that might be involved in extraction of object information are not known. We found that higher visual areas AL, LM and LI had significantly more information about object identity than V1, with area AL consistently outperforming all other areas which is inconsistent with the current assumption that AL is part of a distinct dorsal pathway. Strong interactions between anatomically defined dorsal and ventral pathways in rodents might be particularly important for object detection and discrimination given the importance of navigation in rodents (Froudarakis et al., 2019). To that effect, both areas AL and LI show faster accumulation of information about object identity (**Supp. Fig. 8a**) that could result in the increased temporal stability that has reported recently in higher visual areas of the rat (Piasini et al., 2021). Moreover, the correlations we report in decoding confidence between areas AL and LM (**Fig. 5c**), could be the result of recurrent processes that have been suggested to play a significant role during object recognition (Kar et al., 2019; Tang et al., 2018, 2014; Kar and DiCarlo, 2021). These object-selective dependencies, particularly with area AL showing strong correlations with areas LM and LI, do not share the same structure as have been reported with more parametric stimuli (Smith et al., 2017), which could be due to objects having a statistical structure closer to the preferences of these lateral visual areas. Interestingly, area LI which is believed to be a high visual area did not consistently outperform areas LM and AL and had great variability in the discrimination levels (**Fig. 2d, 3c, 4a**).

Future experiments are required to determine how these different areas work together to extract information about objects that might be used to guide behavior. First, utilizing an ethologically-relevant task while mapping the activities across visual areas might provide even stronger evidence of hierarchy (Hoy et al., 2016). Second, in order to establish a more causal relationship between visual areas and behavior, it will be important to combine behavioral performance with causal manipulations of neural activity. Finally, neural networks models and the inception loop methodology will enable the characterization of the specific features that drive populations of neurons in these different visual areas (Bashivan et al., 2018; Ponce et al., 2019; Walker et al., 2019).

In summary, we offer evidence that mice share similarities with other mammals in their ability to recognize objects. By recording the activity from more than ~400000 neurons across the whole visual system of the mouse, in this paper we have deciphered for the first time for any species how object manifold geometry is transformed to become more separable thus identifying key features of the population code that enable invariant object coding. Given the panoply of tools available, the mouse has the potential to become a powerful model to dissect the circuit mechanisms of object recognition.

## ACKNOWLEDGEMENTS

This work was supported by the the Intelligence Advanced Research Projects Activity (IARPA) via Department of Interior/Interior Business Center (DoI/IBC) contract number D16PC00003 (AST). The U.S. Government is authorized to reproduce and distribute reprints for Governmental purposes notwithstanding any copyright annotation thereon. Disclaimer: The views and conclusions contained herein are those of the authors and should not be interpreted as necessarily representing the official policies or endorsements, either expressed or implied, of IARPA, DoI/IBC, or the U.S. Government. Also supported by R01 EY026927 (AST), NEI/NIH Core Grant for Vision Research (T32-EY-002520-37) and NSF NeuroNex grant 1707400 (AST). HS is partially supported by the Gatsby Charitable Foundation, the Swartz Foundation, the National Institutes of Health (Grant No. 1U19NS104653) and the MAFAT Center for Deep Learning. PB is supported by the the Deutsche Forschungsgemeinschaft (DFG, BE5601/4, SFB 1233 “Robust Vision” 276693517, Cluster of Excellence 2064 “Machine Learning - New Perspectives for Science” 390727645) and the German Ministry for Education and Research (FKZ 01GQ1601, 01IS18039A). EF is supported by a European Research Council (ERC) grant (ERC-2022-STG, NEURACT, Grant agreement No: 101076710).

## AUTHOR CONTRIBUTIONS

**EF**: Conceptualization, Methodology, Validation, Hardware, Software, Data Curation, Formal Analysis, Investigation, Writing - Original Draft, Visualization, Supervision, Project administration; **UC**: Methodology, Validation, Formal Analysis, Writing - Original Draft; **MD**: Investigation, Validation, Formal Analysis, Writing - Review & Editing; **SP**: Methodology, Hardware, Software; **ZT**: Investigation; **TM**: Investigation; **EYW**: Software; **JR**: Methodology, Investigation, Writing - Review & Editing; **PB**: Validation, Writing - Review & Editing; **HS**: Validation, Writing - Review & Editing, Supervision, Funding acquisition; **AST**: Conceptualization, Methodology, Validation, Supervision, Funding acquisition, Writing - Review & Editing

## Materials and Methods

### Stimulus generation

In this study we used four synthesized three-dimensional objects that were rendered in Blender (www.blender.org). Two of the objects were built to match the objects used in (Zoccolan et al., 2009) and the other two were already existing models within Blender. We varied the following parameters of the objects: X and Y location (Translation), magnification (Scale), tilt and axial rotation (Pose) and variation of either the location or energy of 4 light sources (Light). The different object parameters were varied continuously over time in order to generate a cohesive object motion. Objects were rendered either on a gray background, or on a gaussian noise pattern with a fixed seed between objects. The long rendered movie was split into smaller 10 second clips.

### Animals

All procedures were approved by the Institutional Animal Care and Use Committee (IACUC) of Baylor College of Medicine. We used 37 adult mice expressing GCaMP6s in excitatory neurons via either SLC17a7-Cre, Dlx5-Cre, Ai75, Ai148, Ai162 or CamKII-tTA transgenic lines. Nine animals were used for behavioral training (N=9, see **Supp. Table 1**). 31 animals were used in total for two-photon imaging (N=31, see **Supp. Table 1**). Other than a small number of animals that were also used for behavioral training (N=3) the majority of the animals used in imaging were naive (N=28, see **Supp. Table 1**).

#### Behavioral training

The mice (N=9, see **Supp. Table 1**) are trained in a 2 alternative forced choice task in response to moving objects that are presented on a small 7” monitor that is located in front of their home cage. In total, 4 objects are used in the experiments, however it should be noted that 2 objects are presented to each mouse during behavioral training. The training procedure is illustrated in **Fig. 1**. Briefly, naive water restricted mice are placed in a modified cage that has three ports and a monitor on one side of the box. The center port has a proximity sensor, and the two other ports on either side of the central port are used to detect licks and are coupled to a computerized valve-controlled liquid delivery that can deliver liquid volumes with 1uL resolution. The task is as follows: Mice initiate a trial by placing their snout in close proximity to the central port for ~200-500msec. A stimulus that can be one of two objects is presented on the monitor that is ~1.5” in front of the animal. The animal has to report the identity of the object by licking one of the side ports. Each port is allocated to the identity of the same object throughout the training. If the animal licks the correct port, then a small water reward ~5-12*µ*l is delivered almost immediately which the animals consume. A new trial can be started thereafter. If the animal licks the wrong port, a short delay 4-10 seconds is added and the screen turns black. A new trial can start after the delay. Animals have free access to food, and most of the water that they receive comes from their training. The training periods in which animals can initiate tasks are restricted to 4-8 hours a day. At the start of the training animals are shown the same set of clips for each object that contains the same set of transformations. Once animals reach performance levels ( 70% correct), new clips with unique transformations are added. At the end of their training they have seen between 10-20 unique 10s clips of unique object transformations.

#### Behavioral metrics

Lick probability is calculated as the number of trials that the animal responded by licking one port (left or right) divided by the total numbers of trials that the animal gave a response (lick) irrespective of which port. Performance (% correct) is calculated as the number of trials that the animal responded by licking the left port given that left was the correct port plus the number of trials that the animal responded by licking the right port given that right was the correct port divided by the total numbers of trials that the animal gave a response.

#### Generalization test

For the generalization test, at the start of a new session a whole new set of 10 clips are used for each object and the performance was compared to the session that preceded. The pixel model is a linear classifier trained on the pixel intensities of the rendered movies. The Difference of Guassians (DoG) model is used to model the center-surround receptive fields of the retina. The same linear classifier as for the pixel model was trained on the convolution of the rendered movies with the DoG model.

#### Regression analysis

We used a linear regression model to predict the performance of the animals from different object latent parameters: x, y translation offset, magnification, rotation, tilt and speed. Before fitting, the regressors were standardized. We show the total contribution toward explaining the average performance as in (Froudarakis et al., 2014). Error bars shown are the standard errors of the coefficients in the linear regression model scaled by the mean difference delta.

#### Surgery and two-photon imaging

Animals (N=31, see **Supp. Table 1**) were initially anesthetized with Isoflurane (2%) and a ~4mm craniotomy was made over the right visual cortex as previously described (Froudarakis et al., 2014). Animals were allowed to recover for at least 3 days postoperative. On recording day the animals were head-mounted above a cylindrical treadmill and calcium imaging was performed using Chameleon Ti-Sapphire laser (Coherent, Santa Clara, CA) tuned to 920 nm. We recorded calcium traces by using either a large field of view mesoscope (Sofroniew et al., 2016) equipped with a custom objective (0.6 NA, 21mm focal length) with a typical field of view of ~2500×2000*µ*m (N=16), or a two-photon resonant microscope (Thorlabs, Newton, NJ) equipped with a Nikon objective (1.1 NA, 25X) with a typical field of view or ~500×500*µ*m (N=15). Laser power after the objective was kept below ~60mW. We recorded data from depths of 100–380 *µ*m below the cortical surface. Imaging was performed at approximately ~5-12Hz for all scans. During imaging the animals’ eye and treadmill’s movements were also recorded. Scan sessions were continuously monitored by the experimenter to ensure animals were in an attentive state (eyes clear and open) throughout the scan. In cases where the animal was entering inattentive states scan was stopped. Imaging data were motion corrected, automatically segmented and deconvolved using the CNMF algorithm (Pnevmatikakis et al., 2016); cells were further selected by a classifier trained to detect somata based on the segmented cell masks.

#### Receptive field mapping

We mapped the location and size of the RF of the neurons using black and white squares that each covered ~8-10° of the visual field. The squares were presented across the entire range of the monitor in random order for 150-200 ms each. To map the RF of individual neurons we averaged the first 500 ms of the activity of a cell across all repetitions of the stimulus for each location. We fit the resulting 2D map using an elliptic 2D Gaussian. For each neuron we computed the SNR as the ratio of the variance of this image within three SD of the RF center to the variance of the image outside of the three SD of the RF center. In total we recorded 489182 neurons: 222239 in V1, 58159 in RL, 48769 in LM, 24662 in AL, 16681 in PM, 15364 in AM, 11617 in LI, 6845 in LLA, 6231 in P, 6029 in POR, and 72586 neurons not assigned to any visual area.

#### Visual area identification

We generated retinotopic maps of all the visual areas using widefield imaging. The signals from GCaMP6s were captured using either a custom epifluorescence setup or two-photon imaging. For the epifluorescence, brain was illuminated with a high power LED (Thorlabs) and the emitted signal was bandpass filtered at 525nm and captured at a rate of 10 Hz with a CMOS camera (MV1-D1312-160-CL, PhotonFocus, Lachen, Switzerland). For the two-photon retinotopic mapping we sampled the activity from a 2.4×2.4mm area with large field of view two photon microscope (Sofroniew et al., 2016) at a rate of ~5Hz. We stimulated with upward and rightward drifting white bars (speed: 9-18°/sec, width: 10-20°) on black background that had their size and speed constant relative to the mouse perspective as previously described. Additionally, within the bar we had drifting gratings with a direction opposite to the movement of the bar. Images from either the epifluorescent or the two-photon setups were analyzed by a custom-written code in MATLAB to construct the 2D phase maps for the two directions. We used the resulting retinotopic maps to identify the borders and delineate the visual areas as previously described (Garrett et al., 2014) (Wang and Burkhalter, 2007). The cortical spread of all the visual areas exceeded even the coverage from the mesoscope in order to have adequate sampling rate, thus for every animal not all visual areas were imaged. Additionally, the 4mm cortical window was optimized to cover either the lateral visual areas plus V1 or the medial visual areas plus V1.

#### Visual stimulation and reliability

A short 3-5 second segment from 150-380 clips for each object were presented in a random sequence to the left eye with a 25” LCD monitor positioned ~15cm away from the animal. A small number of clips were repeated multiple times in order to estimate the reliability of the neural responses. The reliability was estimated as the correlation of the leave-one-out mean response with the mean response to the remaining trials across repeated movies for each cell and averaged across all cells as previously described (Walker et al., 2019).

#### Discriminability

We used a one-versus-all logistic regression classifier to estimate the decoding error between the neural representations of 2-4 objects of 200-500 ms scenes. Each scene was represented as an N-dimensional vector of neural activity for each response scene. For **Fig. 2** we used N=64 neurons, for **Fig. 3** N=64 and for **Fig. 5** all the recorded neurons from each area, unless otherwise specified. We used a 10-fold cross-validation in which the performance of the decoder was tested on 10% of the data that were held out during training. While in the majority of the animals (N=20/31) we showed the four objects, in some of them (N=11/31) we showed only two objects (**Fig. 2d**, Dots and Diamonds respectively). In order to compare the decoding accuracy of a one-versus-all classifier between experiments that had different number of objects (2 and 4), we converted the decoding error to discriminability, the mutual information (measured in bits) between the true class label *c* and its estimate, by computing

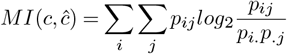

where *p*_*ij*_ is the probability of observing true class *i* and predicted class *j* and *p*_*i*._ and *p*_.*j*_ denote the respective marginal probabilities. For the recordings that we had presented more than two objects, we also computed the pair-wise discriminability for all the pairs of objects and computed the % correct (**Supp. Fig. 3c,d**). For figure **Supp. Fig. 3c** we used all the neurons from each recording.

When generalizing across the background noise in **Fig. 3a-c**, the decoder was trained on 90% of the data with the no-background objects and tested on 10% of the data with the background objects. Instead of the 10-fold cross-validation used in the previous analysis, for the generalization across object parameters in **Fig. 3d,e** we used a 4-fold cross-validation in which the decoder was trained on 75% of the data and tested on 25% with a unique parameter range (out-of-distribution set).

For the low dimensional representation illustrations in **Fig. 2b** and **Fig. 3a** we estimated the distance to the 4 decision boundaries, the linear hyperplane in the N-dimensional neural activity space that separates responses to each object with the rest.

To estimate the discriminability for single neurons in **Fig. 2f** and **Supp. Fig. 3a** we trained the decoder on the activity of each neuron that we sampled for each area and computed an average for each recording recording site.

To estimate the effect of object size and speed in **Supp. Fig. 5**, we binned the object latent parameters and trained and tested the decoder on each separate section of the responses to each bin. Similarly, to estimate how the decoder is affected by the behavioral state in **Supp. Fig. 6**, we estimated discriminability separately for the responses where the animals were running and when they were stationary.

#### Model

For the V1 model, we used the filter responses to a set of 256 localized and oriented filters obtained with Independent Component Analysis (ICA) on 12×12 patches randomly sampled from natural images from the van Hateren database as previously described (Froudarakis et al., 2014) (**Supp. Fig. 4a**). In order to control for the increase in size of the RF in higher visual areas, we also created an enlarged version of these RF (**Supp. Fig. 4a**, ICA 150%) by scaling the initial ICA filters. That method has the disadvantage of changing the spatial tuning properties and thus to create more realistic RF, we linearly combined pairs of the ICA group depending on their response similarity to natural movies. This procedure created larger RF that were more elongated and often had corner-like structure (**Supp. Fig. 4b**, ICA multi). As an additional control, we also used a 144×144 grid of pixels as filters when comparing with the behavioral data (**Fig. 1e**) and 32×32 when comparing to the neural data (**Supp. Fig. 4b, c**). The filter responses were half-wave rectified and squared and were used as a scale to sample from the Weibull distribution with a shape parameter optimized in order to create similar reliability levels to the in-vivo data. Finally, we used the resulting responses to train the same decoder as the in vivo data.

#### Classification capacity and geometry of manifolds

An object manifold is defined by the neuronal population responses to an object under different conditions (i.e. identity-preserving transformations). The ability of a down-stream neuron to perform linear classification of object manifolds depends on the number of objects, denoted *P*, and the number of neurons participating in the representation, denoted *N. Classification capacity* denotes the critical ratio *α*_*c*_ = *P/N*_*c*_ where *N*_*c*_ is the population size required for a binary classification of *P* manifolds to succeed with high probability (Chung et al., 2018). This capacity can be interpreted as the amount of information about object identity coded per neuron in the given population. Capacity *α*_*c*_ depends on the radius of each of the manifolds, denoted *R*_*M*_, representing the overall extent of variability (relative to the distance between manifolds), and their dimension, denoted *D*_*M*_, representing the number of directions in which this variability is spread. These geometric measures are defined through the alignment of the hyperplane (in the representation *N*-dimensional space) that separates positively labeled from negatively labeled manifolds. This hyperplane is uniquely determined by a set of *anchor points*, one from each manifold, that lie exactly on the separating plane. As the classification labels are randomly changed, the identity of the anchor points change; these changes, along with the dependence of the hyperplane orientation on the particular position and orientation of the manifolds, give rise to a statistical distribution of anchor points. Averaging the extent and directional spread of the anchor points with this distribution determines the manifolds radii and dimensions, respectively. Knowledge of manifold radius and dimension is sufficient to predict classification capacity using the relation *α*_*c*_ = *α*_*Balls*_(*R*_*M*_, *D*_*M*_) where *α*_*Balls*_ is a closed-form expression describing capacity of *D*-dimensional balls of radius *R* (Chung et al., 2018).

The separability of manifolds depends not only on their geometries but also on their correlations. For manifold classification with random binary labeling, clustering of the manifolds in the representational space, as expected for real-world object representations, hinders their separability, and the theory of manifold classification has been extended (Cohen et al., 2019) to take these correlations into account in evaluating *α*_*c*_.

Here we used the methods and code from (Cohen et al., 2019) to analyze the geometry of the object manifolds (i.e. manifold radius and dimension) as well as estimate classification capacity of neuronal populations in the different cortical areas. As those methods depend on the correlation structure of the objects, we analyzed neural representations for data-sets of 4 objects (i.e. omitted data-sets where only 2 objects are available). At each session of simultaneously recorded neurons we have sub-sampled from the available population 128 neurons; the subsequent analysis was repeated 10 times with different choices of neurons, and we report the average results across this procedure. Each object manifold is defined by neural responses to an object at non-overlapping 500ms time windows, using the entire range of nuisance parameter space, as well as responses with and without background noise. This analysis was performed at each visual area for sessions where more than 128 neurons are available. The baseline to which classification capacity is compared is the value expected by structure-less manifold which is 2*/M*, where *M* is the number of samples (i.e. time windows where the object was presented).

#### Score Correlations

To evaluate the temporal dynamics and cross-area dependencies of the decoder confidence, we estimated the distance to the decision boundary at different moments within the trial for the object class presented (score values). This decision boundary was a linear hyperplane in the N-dimensional neural activity space (**Fig. 5a**). We then computed the Pearson correlation coefficient between the resulting temporal vectors of the score values across all simultaneously recorded visual areas (**Fig. 5b**, Score correlations). As a control we computed the correlations between the neural responses of all pairs of cells across simultaneously recorded areas (**Fig. 5b**, Pairwise activity correlations). For illustration purposes in **Fig. 5c**, we used the Pearson partial correlation coefficients to account for the co-variability between the areas.

To build a decoder that can decode both the object identity and each of the latent parameters (**Supp Fig. 9**), we split each of the latent parameters distribution into two halves (two groups). We then built a one-versus-all logistic regression classifier to discriminate the neural representations of the 2 objects as well as the group of each of the latent parameter. That resulted in 4 unique classes, e.g. Small Object A vs Small Object B vs Big Object A vs Big Object B. The resulting distance from the hyperplanes was used to compute the score correlations across the different areas. For the control, we randomly sampled from the latent parameter distributions.

#### Temporal dynamics

To evaluate how discriminability evolves across time, we evaluated the performance of the decoder at different time bins after the start of each trial (**Supp Fig. 8**). The trials were split depending on whether the identity of the object of the previous trial was the same (cis trials) or different (trans trials). To compare the performance across the different trial groups we normalized the performances to the average discriminability of the cis trials (**Supp Fig. 8b**) or the trans trials (**Supp Fig. 8c**). To compare across the areas we took the difference of the normalized discriminability with V1 (**Supp Fig. 8d**).

## Supplementary Information

**Supplementary Fig. 1.**
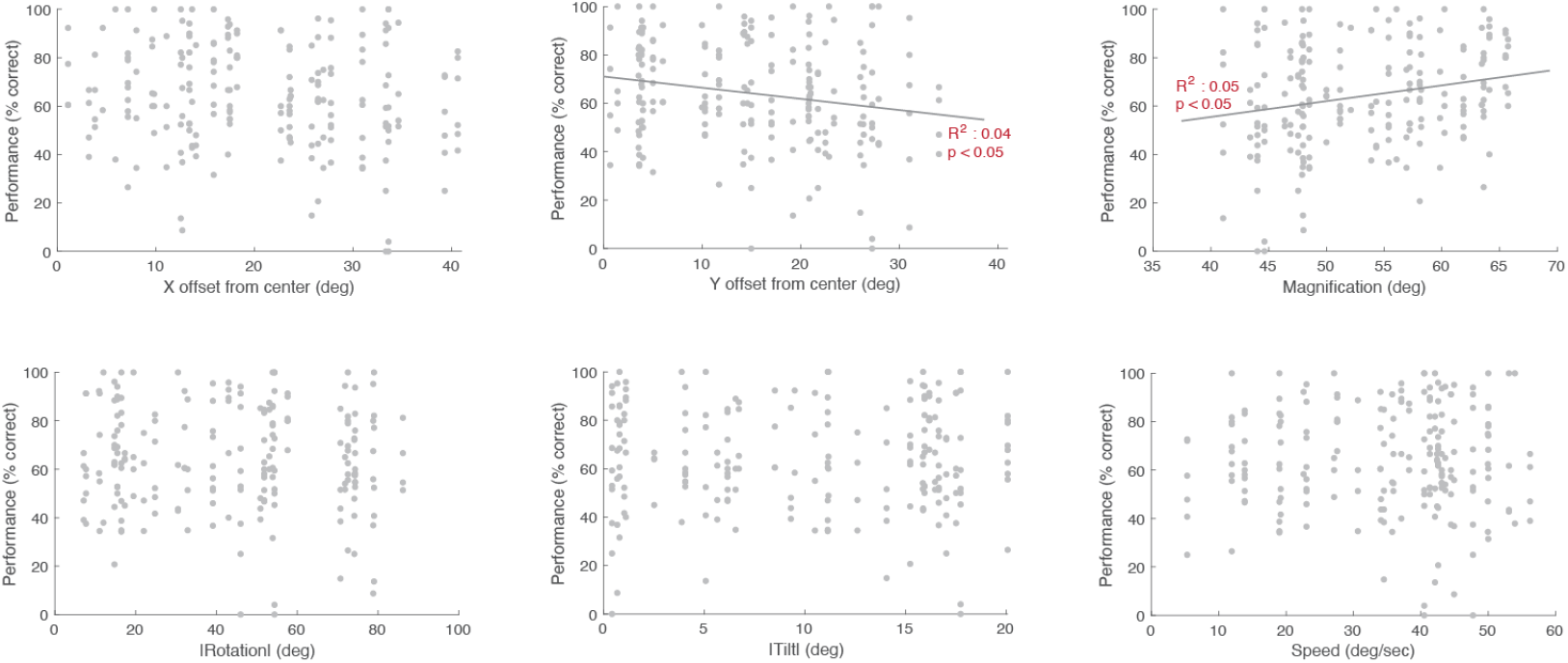
Behavioral performance and object latent parameters. Performance vs object latent parameters: X offset from center, Y offset from center, Magnification, Rotation, Tilt & Speed. X offset, Rotation and Tilt are plotted as the absolute values from the default object parameter set. N=200, all data from 5 animals. The regression line is indicated for each and the explained variance of the regression is noted on the top left of each plot.

**Supplementary Fig. 2.**
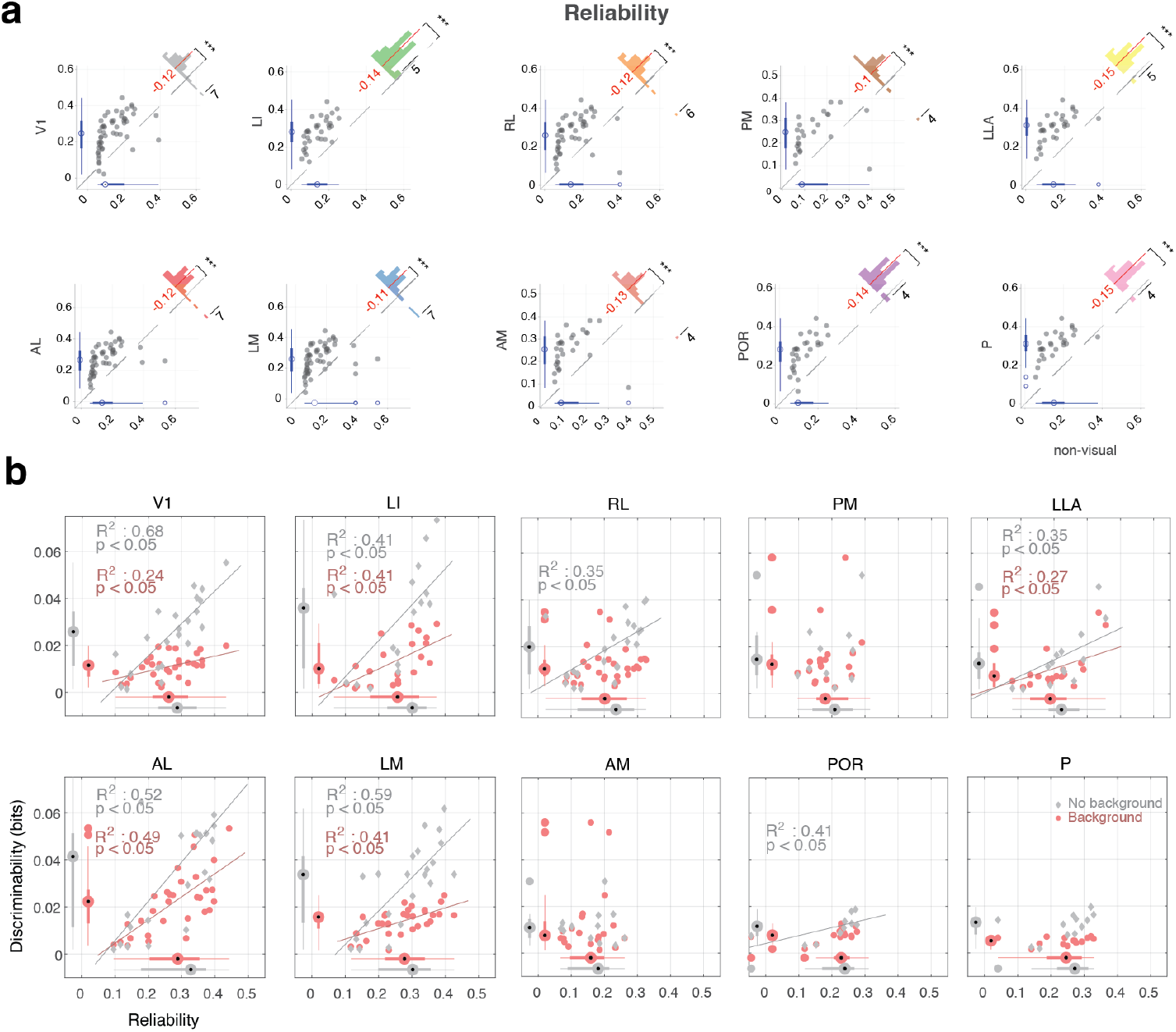
Reliability across all visual areas. **a**. Comparison of the average reliability of the responses to the object stimuli across neurons of all visual areas (y-axis) and neurons in non-visual areas (x-axis). Insert histogram represents the difference between the average reliability between each visual area and non-visual areas. Red line and number indicate the mean difference across all recording sites. Wilcoxon signed rank test *** p < 0.001, ** p <0.01, * p < 0.05. **b**. Discriminability vs average reliability for all the cells with each recording. Plotted separately for objects w/ (red) and w/o background (gray). The regression line is indicated for each and the explained variance of the regression is noted on the top left of each plot. At the sides of each axis are the boxplots for each of the datasets.

**Supplementary Fig. 3.**
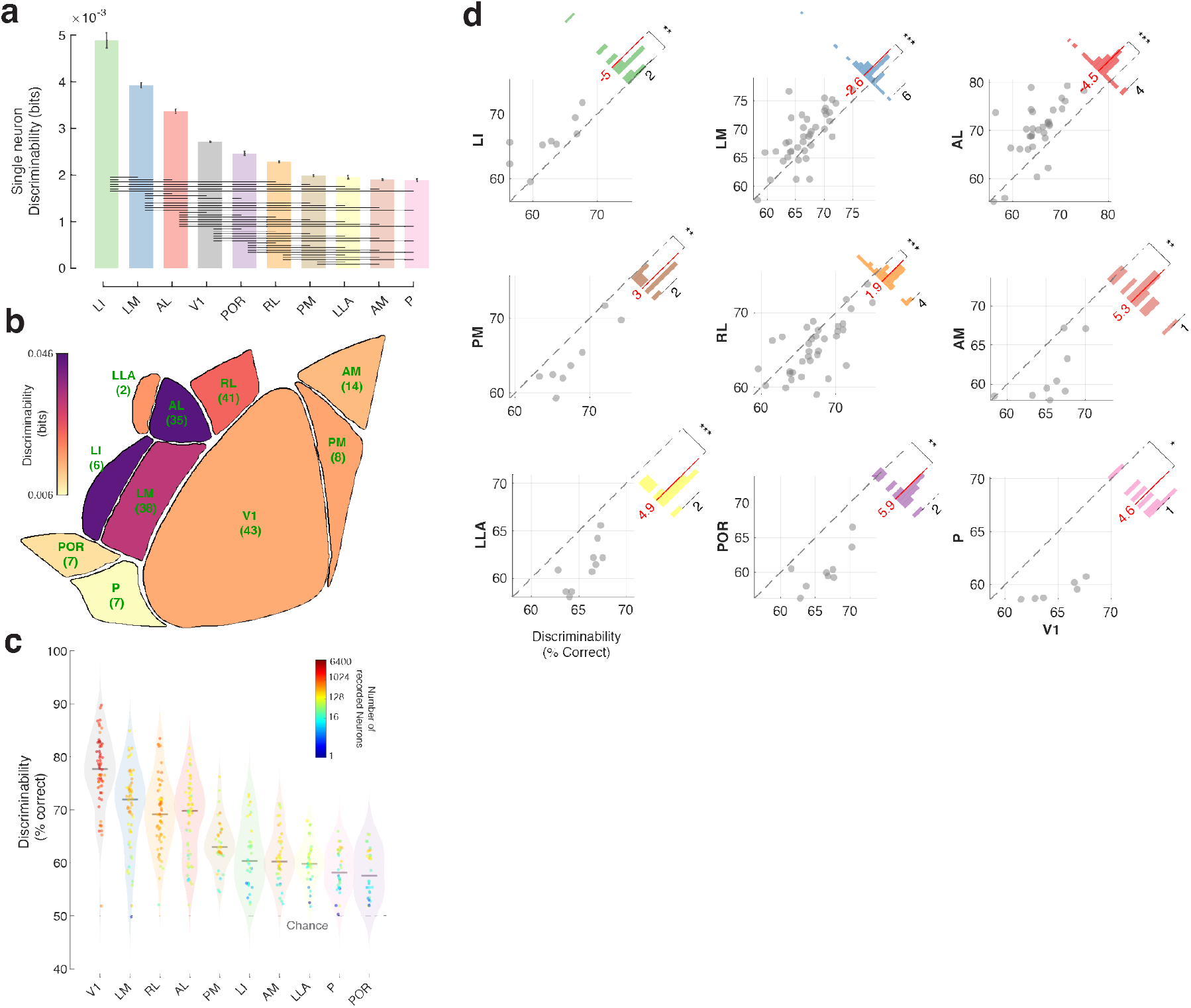
Discriminability with single neurons or with RF restriction or with all the neurons and pairwise decoding of object identity. **a**. Bar graph of the average discriminability when using single neurons to decode the object identity. Horizontal lines indicate p < 0.01 Kruskal-Wallis with multiple comparisons test. Number of recording sites is reported in Fig. 2c. **b**. Average discriminability for all visual areas when selecting a population of 20 cells that have their RF centered within the same 20° of visual space. The number below each area represents the recording sites sampled. * p < 0.05 Wilcoxon signed rank test when compared to V1. **c**. Candle plot of the average pairwise discriminability in % correct. Dots indicate different samples for each area and their color indicates the number of neurons recorded. **d**. Scatter plot of the pairwise object discriminability in % correct of different areas with a population of 64 neurons compared to V1 for all the recording sites. Insert histogram represents the difference between the discriminability of each area and V1. Red line and number indicate the mean difference. Outliers have been omitted for better visualization. Wilcoxon signed rank test *** p < 0.001, ** p <0.01, * p < 0.05.

**Supplementary Fig. 4.**
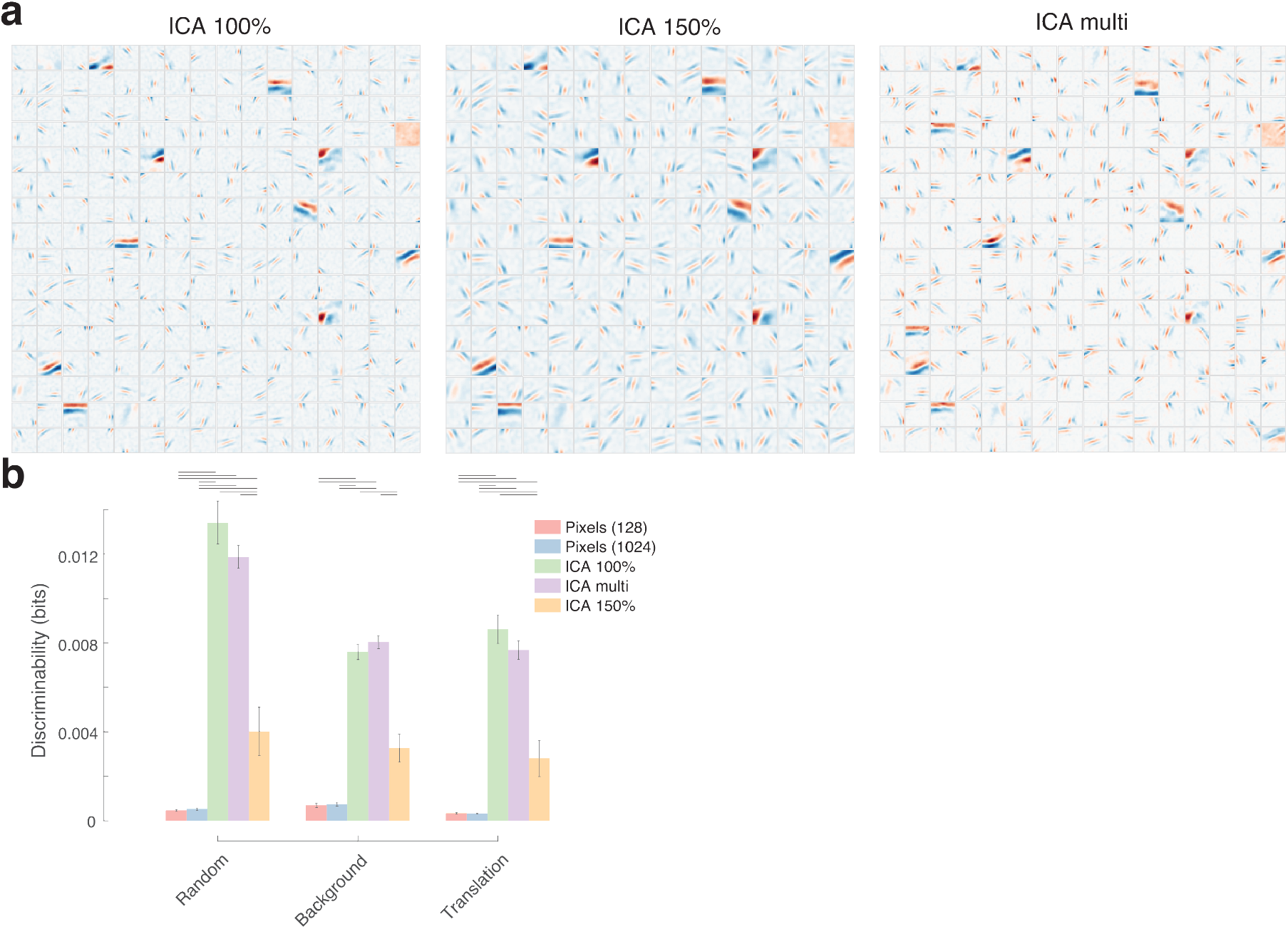
Models of RF and decoding performance. **a**. The 256 RF (ICA 100%), their enlarged version (ICA 150%) and their combination (ICA multi) that were used for the computational model. **b**. Discriminability of the simulated responses of 128 units to objects and the generalization test on objects with background and translation, from the filters in (a) and also the pixels of the stimuli as input. Horizontal lines indicate p < 0.05, Kruskal-wallis multiple comparison test.

**Supplementary Fig. 5.**
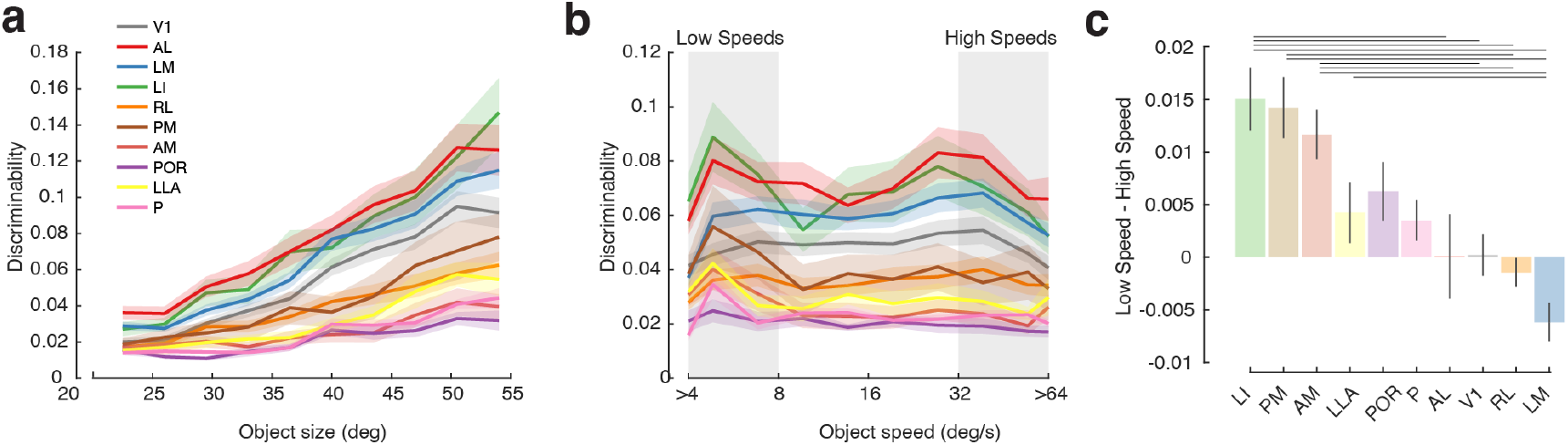
Object size and object speed effect on decoding. **a**. Discriminability as a function of the object size for all visual areas. **b**. Discriminability as a function of object speed for all visual areas. Shaded areas represent S.E.M. **c**. Difference in discriminability between Low (<8°/s) and High (>32°/s) object speeds. Horizontal lines indicate significant difference with Kruskal-Wallis with multiple comparisons test p < 0.05.

**Supplementary Fig. 6.**
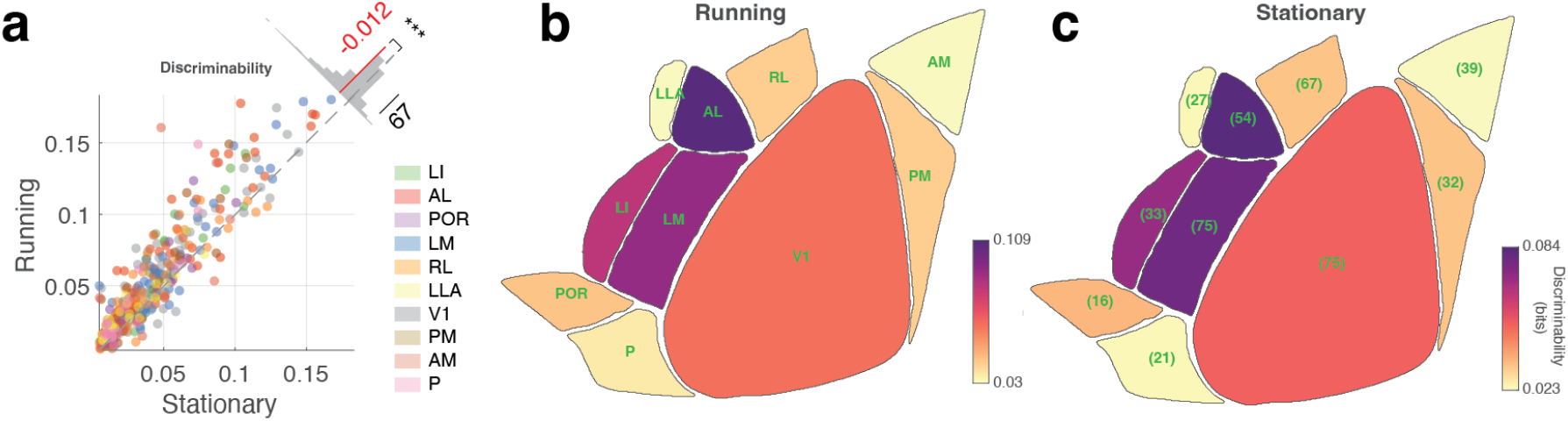
State dependent decoding performance. **a**. Comparison of the discriminability of the responses to the object stimuli across neurons of all visual areas while the animals are running (y-axis) and while the animals remain stationary (x-axis). Insert histogram represents the difference between the discriminability between the two states. Red line and number indicate the mean difference across all recording sites. Wilcoxon signed rank test *** p < 0.001 **b**. Average discriminability for all visual areas while the animals are running and in **c**. while they are stationary. The number in each area in c represents the recording sites sampled.

**Supplementary Fig. 7.**
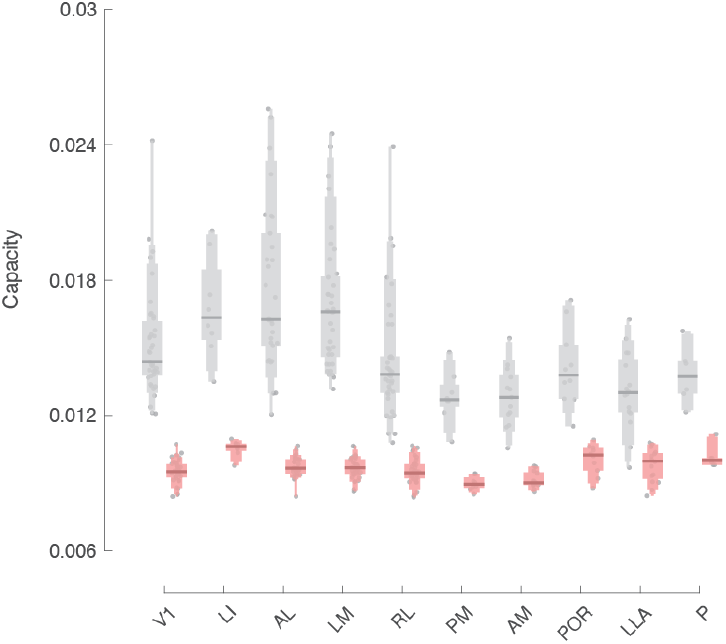
Capacity shuffling control. Gray: Classification capacity of different areas. Red: Classification capacity estimated after shuffling of the object identity labels. Lines indicate the median of the distribution, and the variable bars indicate the 75, 90, 95, and 100 percentiles. Dots represent the raw values.

**Supplementary Fig. 8.**
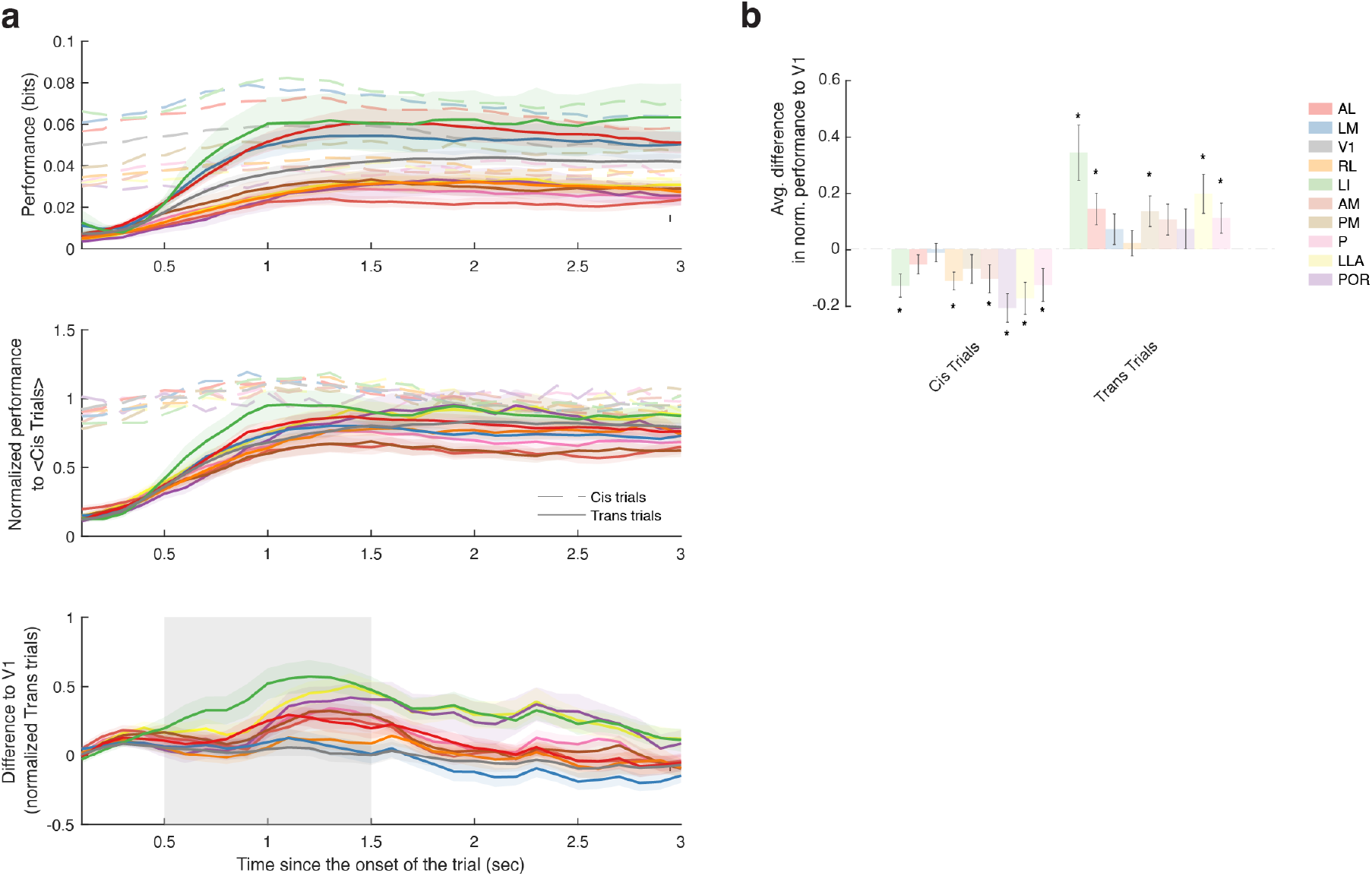
Temporal dynamics when object identity is switching. **a**. Discriminability across time for trials where the preserved object identity is preserved (cis-trials) or switched (trans-trials). **b**. Normalized discriminability to the average discriminability of the cis trials. **c**. Difference between normalized discriminabilities to V1. Shaded areas represent S.E.M. **d**. Bar plot of difference in normalized discriminability of the cis (left) and trans (right) trials during the 0.5-1.5 seconds of the trial. Wilcoxon signed rank test, * p < 0.05.

**Supplementary Fig. 9.**
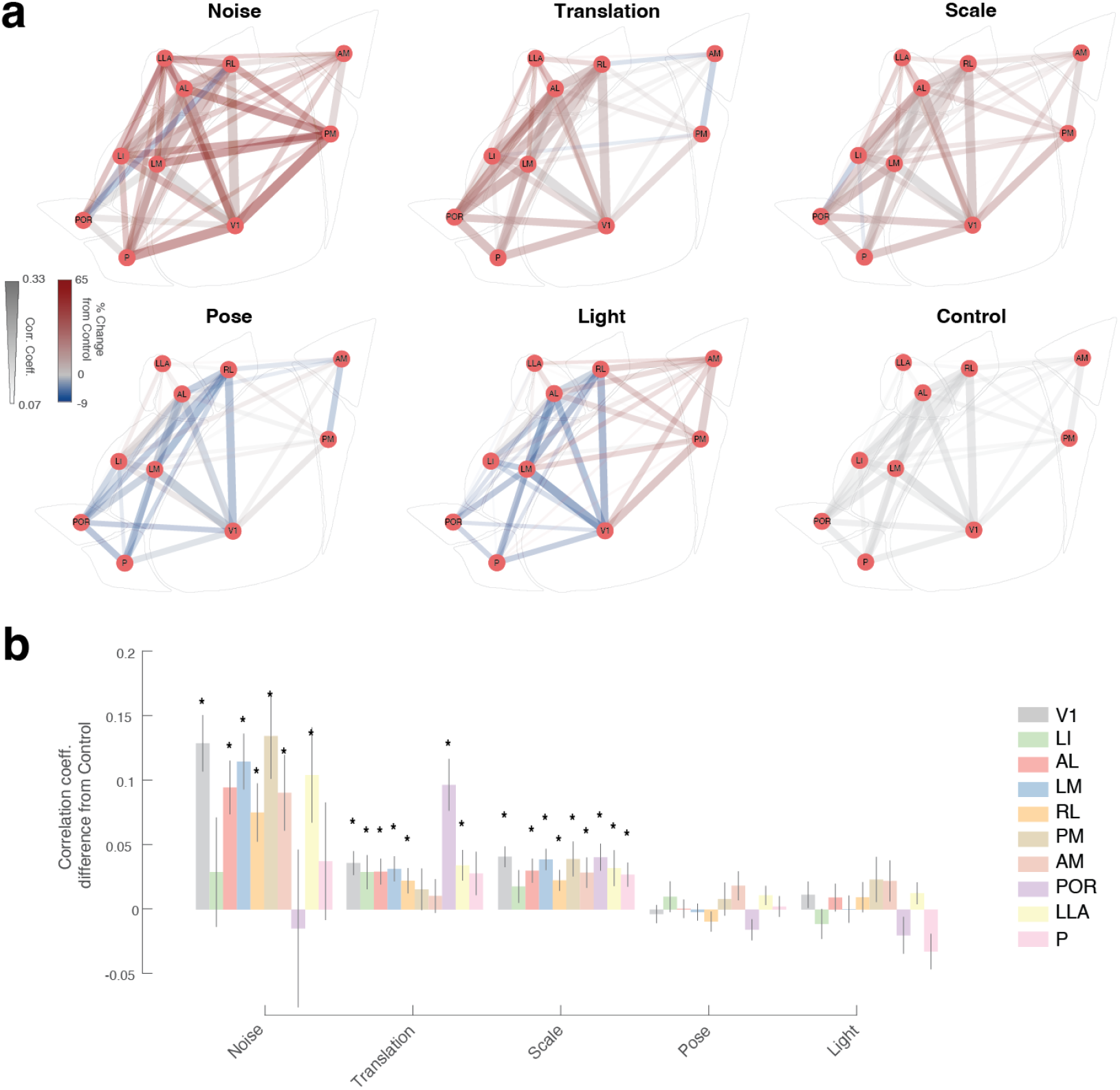
Score correlation for different latent parameters. **a**. Schematic representation of the score correlation coefficients between areas when the the decoder is trained to decode both object identity and the different latent parameters of the objects compared to the control in which we randomly sampled from the latent parameter distributions. The bar width and luminance indicates the score correlation coefficient and the color indicated the % change from the control (bottom row, right) **b**. Average difference between the score correlations of all visual areas to control for different latent parameters, * p < 0.05

**Supplementary Table 1.**
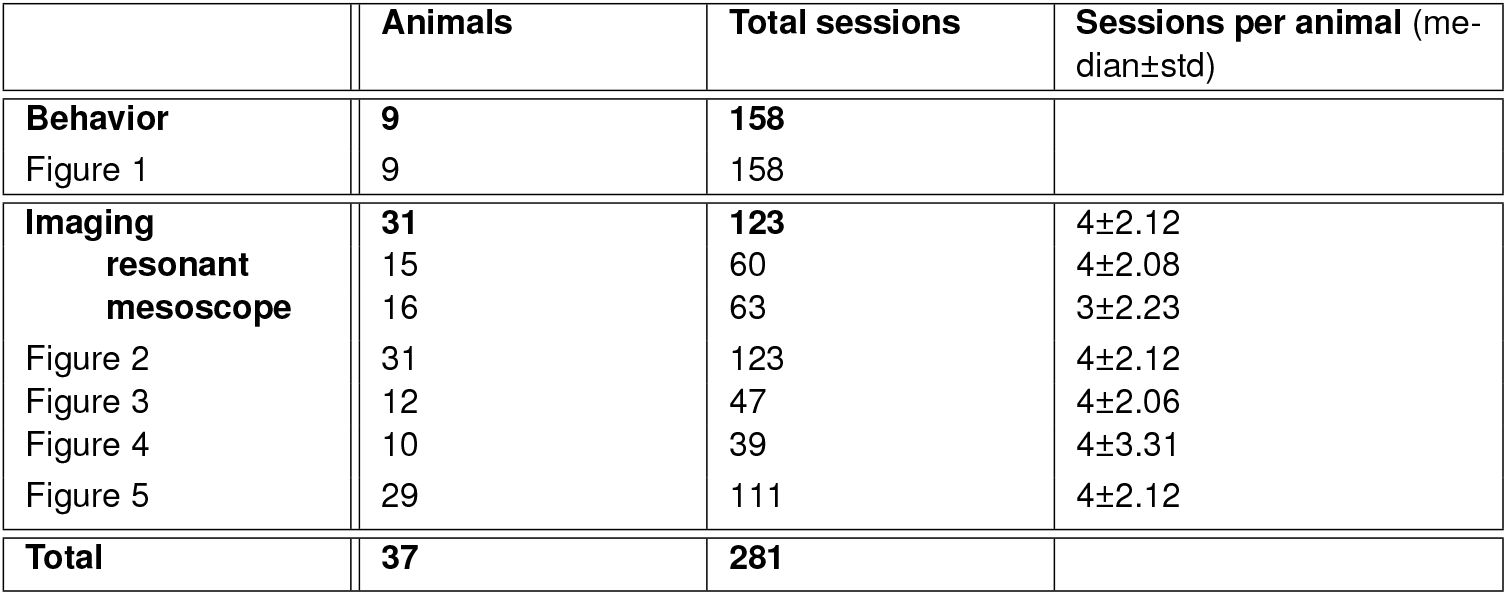
Animals and sessions per experiment type and figure.

**Supplementary Movie 1. Example clips with object transformations used in behavioral tasks**.

**Supplementary Movie 2. Example object clips with objects on a background noise**.

## Notes

### Competing Interest Statement

The authors have declared no competing interest.

### Summary of Updates

Minor modifications in Figures 1,2,3,4,5 supp 8, supp 9 Funding source added

